# Density-dependence and territorial competitors can modulate parrotfish social foraging and herbivory on coral reefs

**DOI:** 10.1101/2025.09.02.673613

**Authors:** Harshul Thareja, Anish Paul, Teresa Alcoverro, Rucha Karkarey, Wenzel Pinto, Siddhi Jaishankar, Siya Bhagat, Rohan Arthur

## Abstract

Social context can modify the behaviour of animals, influencing how they interact with their environment – potentially cascading to effects on ecosystem functions. However, typical proxies for functions, such as biomass or densities, may not completely capture this behavioural variation. We examined how grouping behaviour of an abundant parrotfish species, *Chlorurus sordidus*, influences the critical function of herbivory on coral reefs in the Lakshadweep Archipelago and evaluated the role of density-dependence and territorial aggression from competitors in driving group formation in the entire herbivorous fish assemblage. Feeding rates increased with group size and decreased with aggressive encounters, with parrotfish in larger groups consuming about 80% more algae per capita than solitary individuals. Group foragers also benefitted marginally from reduced territorial aggression from competing herbivores. The propensities of herbivores to form groups, and their group sizes were positively density dependent, and, to a lesser extent, were driven by access to resources defended by territorial competitors. Our study highlights the importance of incorporating behavioural variation in assessments and predictive frameworks of ecosystem functioning. As our results indicate, for social animals, key ecosystem functions are particularly sensitive to animal densities and interspecific interactions, especially when individual function varies with social context.

## 1. Background

Behaviour mediates how animals interact with their environment [1]. These interactions influence ecosystem functioning, or the rates of ecological processes involving energy transfer [2,3]. Since ecosystem functions are aggregates of individual contributions, variations in individual behaviour can scale up to influence ecosystem-scale patterns and processes [3,4]. These influences can be particularly stark for functions contributed by social animals. Coordinated behavioural variations in groups of animals, such as movement and foraging [5], can potentially result in large differences in the distributions of functions across ecosystems [6]. However, conventional methods of quantifying ecosystem functions largely rely on static proxies, such as density or biomass [7], which fail to capture behavioural variation [8], and, consequently, often mischaracterize the rates of ecological functions [9,10]. Similarly, trait-based approaches employing species-averaged trait values are unable to capture differences in trait expression, which may arise from behavioural variation between individuals and populations [11,12].

Recently, research has acknowledged behavioural responses of animals to local cues as a universal modifier of functional roles and expression of functional traits [2,12–14]. For instance, the movement activity levels of fish have been shown to modulate their contribution to nutrient input in estuaries, with more active individuals providing disproportionately more nutrients [3]. Ontogenetic and size associated shifts in the foraging behaviour of a herbivore have been shown to influence the distribution of herbivory in seagrass ecosystems [15]. Thus, explicitly incorporating individual-level behavioural variation within populations in assessments and predictive frameworks of ecosystem functions can lead to more reliable estimates of functional rates [2]. This integration requires (i) finding direct causal links between behaviours and functions [13], (ii) quantifying how variation in these behaviours influences animals’ functional contributions, and (iii) identifying measurable drivers of variation in these behaviours.

Herbivory is a critical function across ecosystems, and variations in herbivore behaviour can strongly modulate the distribution of primary consumption in systems – with far reaching consequences for ecosystem structure and function [16]. Herbivores show considerable variation in foraging associated behaviours, both, within and between species [17,18]. This affects when, where, and how they contribute to primary consumption [8,13]. On coral reefs, herbivorous fish help maintain coral dominance by mediating the relationship between coral and algae [19,20]. This function is critical to reef resilience under climate change, as reefs are subject to increasingly frequent coral mass mortalities. [21]. Many herbivorous fish typically forage in fission-fusion groups, which dynamically vary in their size and composition [22–24]. Foraging behaviours are often modulated by an animal’s social environment, and these foraging decisions could therefore have important consequences for their contribution to herbivory [17,25]. For instance, group sizes can influence home range sizes, changing where an animal feeds, and resource intake rates, affecting the quantities it consumes [17,26]. This variation in the social environment of individuals can be driven by dynamic trade-offs between the costs and benefits of grouping, which vary in response to the abiotic environment and biotic interactions within and outside the group [27].

Animals form groups for benefits associated with foraging, predation avoidance, and reproduction ̉[28,29]. Several hypotheses have been proposed to explain group formation in herbivorous fish, but their relative importance remains unresolved [30,31]. One prevalent hypothesis suggests that grouping is driven by interspecific interference competition for a resource– roving herbivores form groups to access resources (algae) defended by benthic territorial fish which guard this resource (henceforth, resource-competition hypothesis) [22]. While group formation enables individuals to overcome the territorial defence of algae-farming damselfish[32–34], it is not clear to what extent this benefit drives grouping behaviour. Additionally, a large part of the benthos on coral reefs is often defended by non-farming territorial fish, such as surgeonfish (Acanthuridae), and whether the resource competition hypothesis can be more generally applied to all territory-holding species – which may defend a lower quality but more widespread resource than damselfish – remains untested. Another critical untested prediction of this hypothesis is that an increase in the proportion of defended resources, or stronger resource defence, should lead to an increase in grouping tendencies and group sizes of roving herbivores. Alternatively, grouping behaviour could be density dependent [35]. At higher densities of potential group participants, the probability of encountering potential partners increases, and individuals may have a larger pool from which to select group partners, thereby minimizing the activity-matching costs of grouping [36]. Evaluating these hypotheses and estimating the relative role of hypothesized drivers can help us predict how grouping behaviour and the function of herbivory could respond to altered species interactions and community composition. While individual-level investigation is useful for identifying mechanisms, coupling it with assemblage-scale investigations can give insights into whether the mechanisms generalize across the assemblage and give rise to expected patterns at functionally relevant scales.

Here, we link behavioural variation (group size) to an ecosystem function (herbivory) and identify drivers of variation in this grouping behaviour (figure 3). For the first objective, we examine the influence of grouping behaviour on an individual’s contribution to herbivory. We then, for the second objective, test two potential drivers of variation in the grouping behaviour of herbivorous fish – access to resources defended by territorial benthic competitors and herbivore densities. Specifically, for the first objective, we asked how group sizes influence the feeding rates of individual *Chlorurus sordidus*, an abundant coral reef herbivorous fish. We expected individuals in larger groups to have higher feeding rates if they were participating for foraging benefits [29]. Then, we tested two behavioural predictions of the resource competition hypothesis – *C. sordidus* individuals receiving more aggression from territorial competitors should have lower feeding rates, and *C. sordidus* in larger groups should benefit by receiving less aggression from territorial competitors. For the second objective, we scaled up to the entire herbivore assemblage and used community censuses to quantify the influence of the densities of (i) territorial competitors and (ii) potential group participants (herbivores) on the grouping propensities and mean group sizes of roving herbivorous fish.

## 2. Methods

### (a) Study area

We sampled reefs at two coral atolls, Kadmat and Kavaratti, in the Union Territory of Lakshadweep, India (8° N – 12°N, and 91° E – 74° E) which forms the northern end of the Chagos-Lakshadweep volcanic ridge (figure S1). These reefs host a diverse assemblage of herbivorous fish, which shows spatial variation in structure and function [37]. Grouping behaviours in this assemblage are well-described, and these islands are among the few locations where grouping behaviour of reef fish has been systematically studied [23,24].

### (b) Study system

In this region, the herbivorous fish assemblage that feeds on epilithic algal matrix (EAM) consists mainly of species from Acanthuridae (surgeonfish) and Scarini (parrotfish) [37]. Species like *Acanthurus lineatus*, *Acanthurus leucosternon*, and *Ctenochaetus striatus* (a detritivore), typically hold territories (henceforth territorial competitors). In contrast, Scarini (dominated by species like *Chlorurus sordidus*, *Scarus psittacus*, *Scarus prasiognathus*, *Scarus rubroviolaceus*, and *Chlorurus strongylocephalus*) and some Acanthurids (like *Acanthurus auranticavus* and *Acanthurus nigricauda*) are typically roving herbivores which try to access algal resources over larger areas, including within the territories of the benthic territorial competitors, and often form foraging groups [23].

We used a combination of focal behavioural observations and herbivore assemblage censuses to evaluate the factors influencing feeding rates and grouping behaviour of herbivores (figure 3, table S1). We selected *Chlorurus sordidus*, a small excavating parrotfish [38] for behavioural observations because it was the most abundant and most commonly grouping parrotfish in our study sites and showed high plasticity in group sizes and composition. All data was collected using SCUBA between January and May 2024.

### (c) Focal behavioural observations

At six sites on two coral atolls – Kadmat and Kavaratti *(figure S1)*, we conducted 103 focal behavioural observations [39] of individuals of the parrotfish *Chlorurus sordidus* foraging in a range of group sizes (from solitary to up to 32 individuals) (Table S1; total time = 341 minutes). This included monospecific groups (n = 65), and multi-species groups (n = 27) groups (defined as any group with more than one species). All observations were conducted at depths of 6-12 m. We defined a foraging group as a congregation of fish that moved together or, in cases where the group did not move, a congregation that foraged together in the same area for the entire length of the observation period, and where individuals were not separated by more than 10 body lengths [23,24]. While some studies have used shorter inter-individual distances to define cohesive groups of fish, these were not suitable for our study because herbivorous fish form dynamic fission-fusion groups, and 10 body lengths falls within the range used by previous studies of mixed-species groups [40]. On encountering a group, we haphazardly selected one individual between 12-18 cm total length for our behavioural observations. We followed it until the group dissipated, or for five minutes, whichever was sooner. We discarded observations where the group dissipated in less than 90 seconds, or if individuals showed signs of being disturbed by our presence (accelerating away from the observer or repeated hiding under reef structure) [39]. We maintained a minimum of two-meter distance from the group which, based on our preliminary observations, did not appear to perceivably influence the focal individual. On the rare occasion when we lost track of the focal individual, we immediately switched to another individual of the same body size in the group, in the same behavioural state, and in the same part of the group [32,33]. If this was not possible, we abandoned the observation. The same area was not sampled twice to avoid repeat observations of the same groups or individuals.

During each observation, one observer noted the bites per feeding bout of the focal individual, the group composition (count, body length, and species identity of each group participant), and the species identity of the benthic territorial competitor for each aggression event received by the focal individual. Since foraging groups showed a high degree of fission-fusion dynamics, we considered an individual to be part of a group if it was present in the group for more than half the observation period. Simultaneously, the second observer conducted one-to-three point counts (depending on the distance the group covered during the observation) within a three meter radius centred around the area where the focal individual was feeding to note the abundance of territorial damselfish and surgeonfish. If the group did not move beyond the three-meter radius, the count was not repeated. The second observer also visually estimated the percentage EAM cover, and structural complexity (on a scale of zero, completely flat, to five, canopy height > 1.5 m) within the same three-meter radius [41]. We divided total bites by the duration of an observation to calculate bite rates per minute, a metric of resource acquisition rates. We restricted our behavioural observations to mornings (9 AM to 12 PM) in Kadmat and evenings (3:30 PM to 6:00 PM) in Kavaratti due to logistical constraints during Ramadan, but all observations within a site were conducted in the same time window. To account for differences in sampling time between sites, we included site as a grouping variable with a varying intercept in our multilevel models.

### (d) Quantifying fish assemblage composition, grouping propensities, and group sizes

We sampled the fish assemblage at 15 sites across Kadmat and Kavaratti atolls *(figure S1)*. We conducted this sampling independent of the focal follows on separate sampling instances. Since the benthic composition and herbivore assemblage varies between the eastern and western aspect of the islands we sampled, we selected sites to maximise spatial coverage on either aspect of each island and to capture wide gradients of herbivore densities [37,42]. We sampled each site using haphazardly laid underwater visual censuses based on timed swims (8.5 minutes) along the depth contour, each covering approximately 70 m (table S1). Swims were conducted at a slow constant pace to minimize variations in distance travelled. We divided our sampling effort into two depth classes – 9-14 meters and 5-8 meters, and sampled 2-4 timed swims per depth class within each site (n = 73 timed swims).

On each timed swim, one observer noted the counts of herbivores (Acanthuridae and Scarini) as solitary or grouping individuals, the size of groups, and the species identity and total length of all herbivores larger than 10 cm encountered within a 10m belt. This was used to calculate the grouping propensity at each transect (proportion of the total herbivorous fish forming groups). The composition of each group was also noted. While previous work has used timed random swims to sample multi-species groups paired with separate transects to sample the herbivore assemblage [23,24], we used linear timed swims to ensure that group composition and the herbivore assemblage were sampled at the same spatial scale in a paired manner, while also being similar to standard belt transects widely used to sample fish communities on coral reefs. The second observer noted the species identity and total length of all piscivorous fish (> 35 cm). We classified fish into functional guilds based on FishBase [43]. In addition, at three points along each transect (at the start, middle, and end), one observer noted the species identity, total length, and counts of territorial damselfish (Pomacentridae) in a five-meter radius, and visually estimated percentage EAM cover and structural complexity. All data was collected by the same two observers.

### (e) Graphical Causal Models (GCMs) and data analyses

We used the graphical/structural causal modelling framework [44] to investigate the relative importance of factors influencing individual foraging rates of the parrotfish species *Chlorurus sordidus* and grouping behaviour of herbivorous fish in coral reefs. This causal inference method uses Directed Acyclic Graphs (DAGs) to represent the causal knowledge and assumptions about a system, and provides a set of rules to guide statistical analysis and arrive at causal effects of interest while blocking non-causal biasing paths. Our first DAG represents factors influencing foraging rates of individuals of the species Chlorurus sordidus (figure 1a), and the second DAG represents factors influencing the grouping tendencies and group sizes of herbivorous fish at a site (figure 2a), including confounders and mediators. In both the DAGs, “site” was added post-hoc to represent potential unidentified or unmeasured site-level confounders [45]. Details about the structure, construction, and implementation of the DAGs is provided in the electronic supplementary material (ESM) (Section 1, Tables S2-S3). Interpreting our results as causal effects requires the assumption that there are no confounding variables other than those specified in the DAGs.

**Figure 1.**
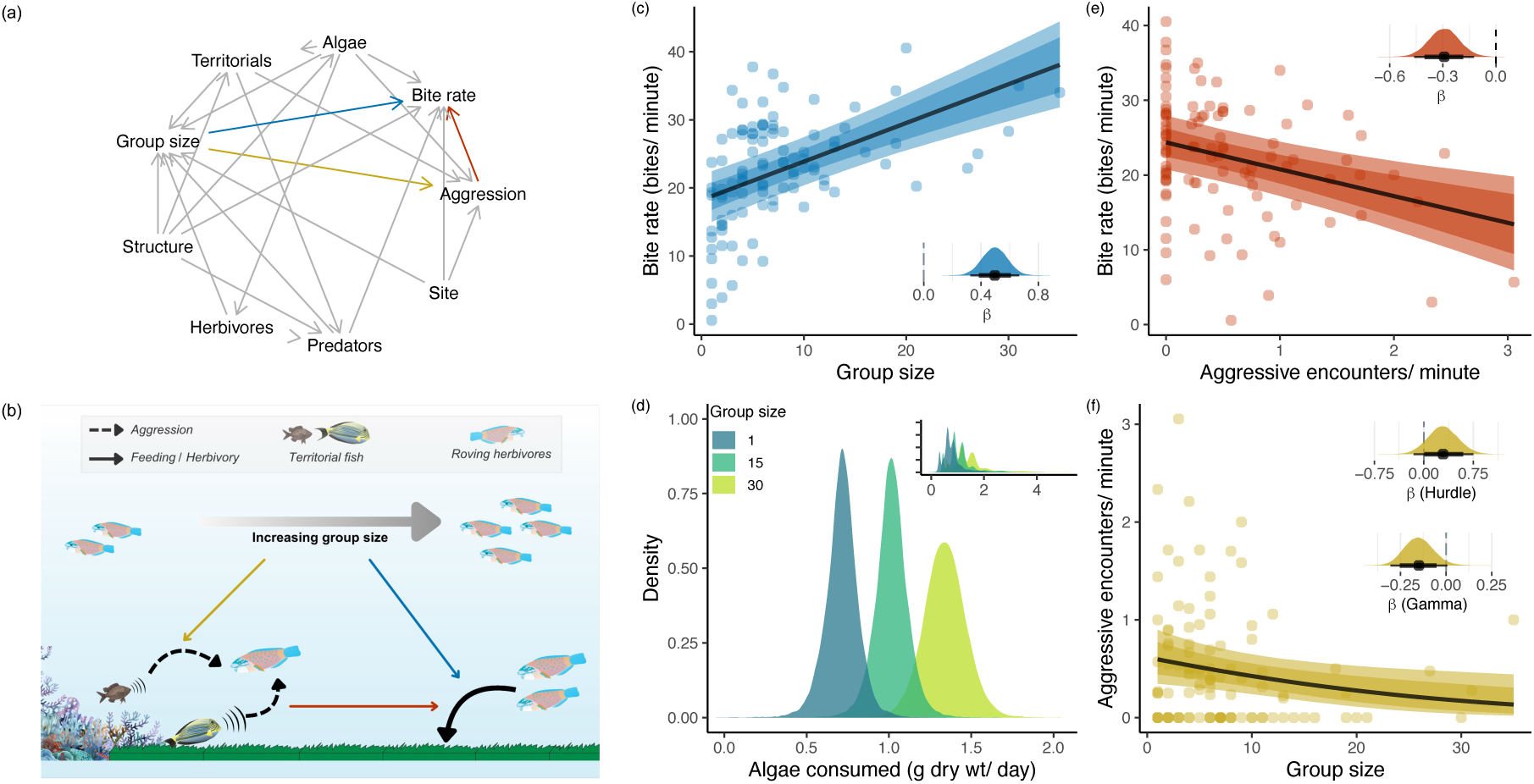
Social foraging facilitates herbivory, but the foraging benefit is only weakly mediated by dilution in aggression. **(a)** Directed Acyclic Graph (DAG) representing foraging by individual *Chlorurus sordidus*. Each directed arrow represents a hypothesised causal relationship, and the lack of an arrow conveys an assumed lack of a direct causal relationship. **(b)** Conceptual diagram summarising the behavioural relationships investigated for *C. sordidus*. Coloured arrows correspond to relationships sharing the same colours in other panels. We investigated the effects of group size on individual feeding rates (blue) and the rates of aggression received from territorial competitors (yellow); and the effect of aggression received on feeding rates (red). **(c)** Group size had a direct positive effect on feeding rates of *C. sordidus,* providing a foraging benefit in larger groups; **(d)** expected marginal predictions of potential algal consumption by a *C. sordidus* individual feeding alone and in groups of 15 or 30 individuals assuming a body size of 14.5 cm (main figure) and using the observed distribution of body sizes (inset); **(e)** shows the direct negative effect of aggression received from territorial competitors on bite rates suggesting that receiving aggression could be a cost to foraging, and **(f)** shows a minimal negative effect of group size on aggression received by *C. sordidus*. **(c)**, **(e)**, and **(f)** show conditional effects from corresponding models with other variables set at their mean; shaded regions represent 80% and 95% credible intervals around the posterior mean (black line); points are raw data. Inserts are posterior distributions with mean and credible intervals (circle and line) without back-transformation. Fish vectors are from the Integration and Application Network, University of Maryland Media Library.

**Figure 2.**
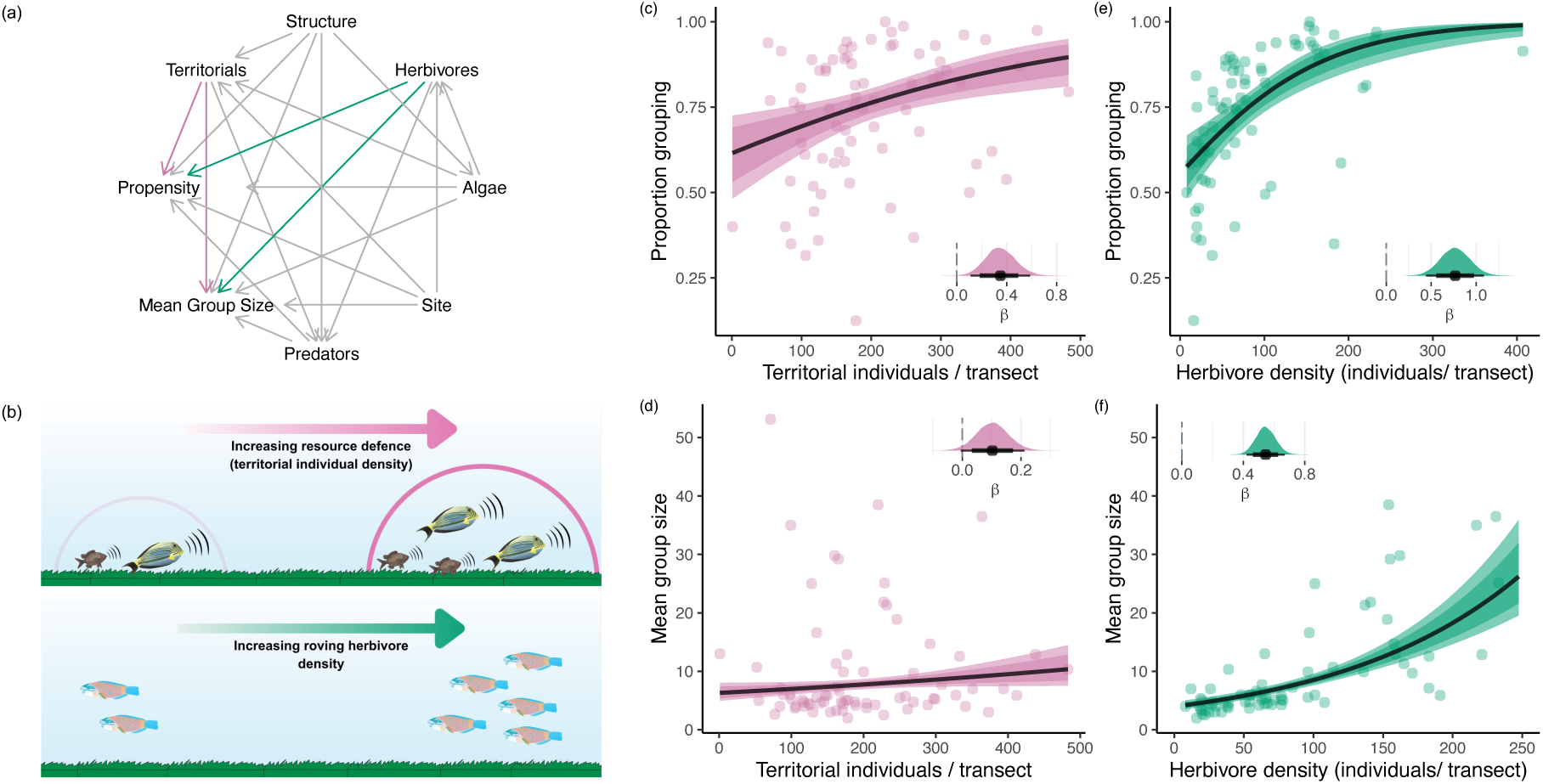
Influence of resource defence by territorial competitors and density dependence on grouping propensities and mean group sizes of grazing herbivorous fish. **(a)** Directed acyclic graph for factors influencing grouping propensity and mean group sizes of a herbivorous coral reef fish assemblage, coloured arrows represent estimated relationships of interest and correspond to colours in other panels. **(b)** Illustration showing gradients of the density of territorial competitors (resource defence) and grazing herbivores. **(c)** to **(f)**: Graphs showing the direct effects of density of territorial individuals on grouping propensity **(c)** and mean group sizes **(d)**; and the direct effects of grazing herbivore density on grouping propensity **(e)** and mean group sizes **(f)**. **(c)** to **(f)** show conditional effects from corresponding models with other variables set at their mean; shaded regions represent 80% and 95% credible intervals around the posterior mean (black line); points are raw data. Inserts are posterior distributions with mean and credible intervals (circle and line) without back-transformation. The x-axis in **(f)** was truncated at 250 for clear visualisation. Fish vectors are from the Integration and Application Network, University of Maryland Media Library.

**Figure 3.**
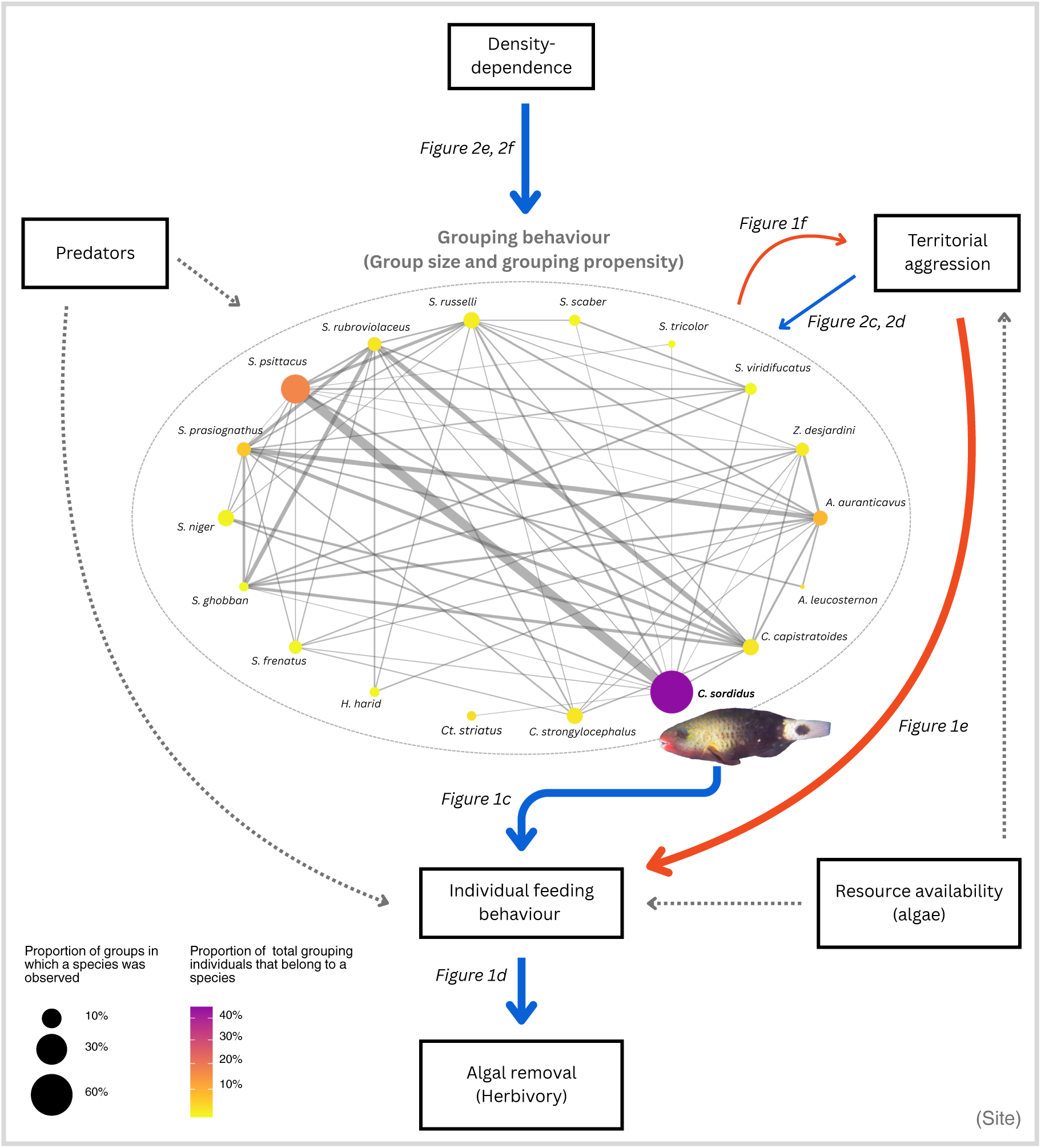
A schematic of the quantified effects and network plot summarising group composition observed in 560 multi-species herbivore groups across 73 timed swims. Solid arrows represent effects of interest (blue – positive effects; red – negative effects) and dotted arrows represent the effects that were controlled for. Thin coloured arrows represent weak effects. Figure numbers corresponding to the effects of interest are indicated. In the network plot, each node represents a species. The size of a node represents the proportion of groups in which a species was observed. Node colour represents the proportion of total grouping individuals across all surveys of that species. Edge width represents a measure of association strength – simple ratio index (SRI) = AB / A + B + AB, where AB is the number of times both species occurred in groups, and A and B are number of times one of the species occurred in groups [56]. Three species which were only observed in single species groups are not presented. (Abbreviations – *C.: Chlorurus; S.: Scarus; Ct.: Ctenochaetus; H.: Hipposcarus; A.: Acanthurus; Z.: Zebrasoma*). *C. sordidus* picture: Rickard Zerpe, CC BY 2.0 (Wikimedia Commons)

We tested our DAGs for consistency with data (ESM S2, Tables S4-S5), and subsequently used them to select adjustment variables for each direct relationship of interest such that all non-causal and indirect paths were blocked [46] (ESM S3). To account for potential unobserved and unidentified confounding variables at the site level we used a group mean covariate design in our statistical models, a type of correlated random effects design [45] (ESM S1 and S3). We calculated group level, or site-level, means of a causal variable of interest and included them as a predictor. We used site as a random intercept to account for the clustering of observations.

#### (i) Effect of group sizes on herbivory

To quantify the effect of group sizes on herbivory, we estimated the direct effect of group sizes on the bite rates of *C. sordidus* (figures 1a, 1b, 3). The statistical model (with scaled predictors and response) derived from our DAG (ESM S3) was :

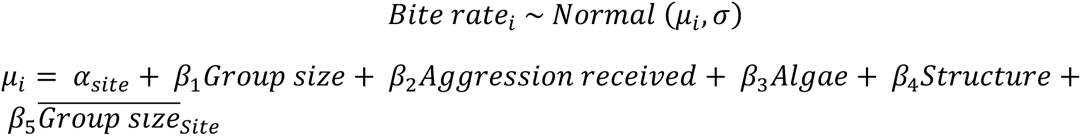

We used a normal distribution because the coefficients were easy to interpret, and the model had an adequate fit.

Then, to estimate per-capita herbivory differences for an individual feeding in groups of different sizes, we first calculated the biomass of algae removed per bite by an individual using a published relationship between bite size and body length for Scarids [47,48]:

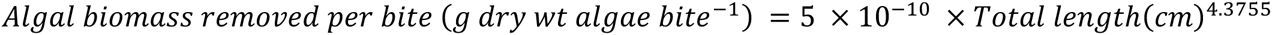

We did this (i) for individuals of 14.5 cm because that was the mean total length of individuals in our observations, and (ii) using the observed distribution of body sizes from the behavioural observations. Then, assuming 10.5 hours of foraging per day, we used our statistical model of bite rates to predict the total bites taken by an individual in groups of different sizes. We multiplied total bites per day with amount of algae removed per bite to calculate the biomass of algae removed per day, for individuals feeding in groups of different sizes. The relationship used to estimate algal biomass removal potential was derived for Caribbean parrotfish species [47,48], thus we interpret the relative differences in estimated values in groups of different sizes instead of absolute values.

#### (ii) Testing the resource competition hypothesis at the individual-scale

First, to test whether receiving aggression from territorial competitors could be a foraging cost, we estimated the effect of aggression from territorial competitors on the feeding rates of *C. sordidus* (figure 2a, 2b). According to our DAG, our previous model of bite rates was consistent with inferring the effect of aggression on bite rates since it blocked all biasing paths by controlling for the confounders – algae, group size, structure, and site. Then, to test a prediction of the resource competition hypothesis – that individuals in larger groups should benefit by receiving lower aggression from territorial competitors, we wanted to estimate the effect of group sizes on aggression rates received by herbivores. Since many individuals did not receive any aggression, we used a hurdle-gamma model for aggression rates. A hurdle model is a two-part model – the first part models whether the response (aggression rate) is zero or non-zero using a logistic model, and the second part models the non-zero data as another distribution [49,50]. While hurdle models are generally used to model two step processes [51], we used the first part of the hurdle-gamma model – the logistic part – to interpret whether group size influences the probability of receiving no aggression, and the second part – the gamma model – to estimate how group size influences aggression rates for individuals that do receive aggression. We removed a group of 26 individuals from this analysis because our focal individual in that group received more than six aggressive encounters from the same *A. lineatus* individual towards the end of our observation, was chased for a very long distance, and then left the group. The model structure (with scaled predictors) based on our DAG, which controlled for the two confounders, algae, and total aggressors in the feeding area, was:

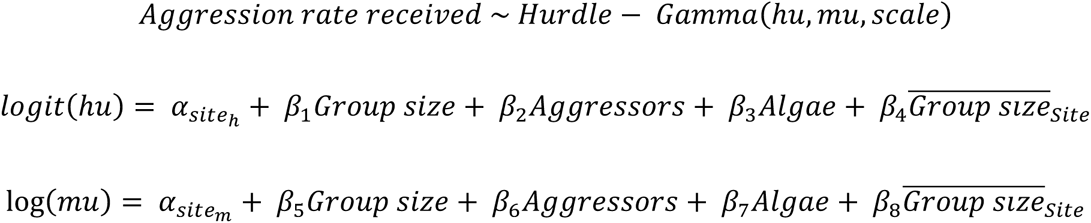

#### (iii) Community-scale drivers of herbivore grouping

We tested how the extent of resource defence (densities of territorial competitors) and herbivore densities influence the grouping propensities and mean group sizes of herbivores. We modelled the grouping propensities of herbivores, or proportion of the herbivore assemblage that was participating in groups, on a transect using a beta multilevel model. We considered *Acanthurus leucosternon*, *Acanthurus lineatus*, *Ctenochaetus striatus, Plectroglyphidodon lacrymatus,* and *Pomacentrus chrysurus* to be territorial herbivores based on our observations and published evidence that they hold territories and aggressively chase intruders [17,52]. On one transect, we encountered a group of *A. leucosternon*. We did not include these individuals in either – group or territorial individuals. We jittered one response from a value of 1 to 0.99999 to be able to model the proportion of shoaling individuals as a beta distribution. The model structure (with scaled predictors), consistent with inferring the effect of both herbivore density and territorial competitor density (ESM S3) was:

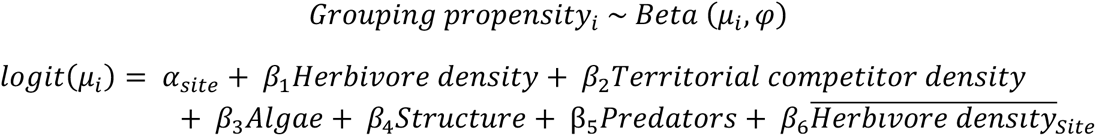

For modelling mean group sizes on a transect, we used a gamma multilevel model. The model structure (with scaled predictors) (ESM S3) was:

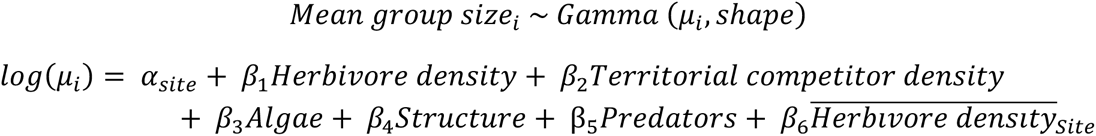

##### Model fitting

We fit all Bayesian models using the R (v. 4.4.1) [53] package *brms* [54], which uses STAN. We ran our models with four Markov chains, having 8,000 iterations each of which 1,000 were warmup iterations. We increased the adapt delta control parameter to reduce the step size and reduce divergent transitions in the MCMC sampling to under three. We visually assessed model fit using posterior predictive checks, and checked for convergence using trace plots, effective sample sizes, and R-hat values [55]. We used informative priors suggestive of the prior expectations of directions of effects, but wide enough to not constrain posteriors to either direction. Rationale for the prior distributions, which were decided based on literature and domain knowledge is provided in the ESM (tables S6 – S9).

To test the sensitivity of our inferences to certain assumptions, exclusion of some data, and choice of priors, we performed a series of sensitivity analyses (ESM S5).

## 3. Results

### (a) Per-capita herbivory increases with group sizes

Increasing group sizes had a direct positive effect on the bite rates of *C. sordidus* (β = 0.50, 95% credible interval (CI) = 0.33 to 0.66; figure 2c; table S9), indicating a foraging benefit in larger groups. This translated to differences in functional contributions in the form of higher per-capita algal consumption in larger groups, with individuals in groups of 30 having almost 80% higher algal consumption potential compared to those feeding alone (figure 1d).

### (b) Access to defended resources may weakly mediate individual benefits of social foraging

Aggression from territorial competitors reduced a forager’s feeding rates (β = -0.29, 95% CI = -0.46 to -0.12; figure 1e; table S10). However, increasing group sizes led to only very weak reductions in per-capita aggression received from territorial competitors (figure 1f). The logistic part of the hurdle gamma model suggested negligible changes in the probability of receiving zero aggression in different group sizes (β = 0.29, 95% CI = -0.15 to 0.75, logit scale; table S11) since the posterior distribution of the hurdle parameter was centred close to zero. Meanwhile, the gamma part suggested that increasing group sizes led to a weak reduction in aggression received by a forager – with ∼97% of the posterior probability of the group size coefficient lying below zero (β = -0.15, 95% CI = -0.30 to 0.01, log scale; figure 1f). However, the strength of this effect was also very weak.

### (c) Group composition and drivers of grouping for the herbivore assemblage

We recorded 21 species of grazing herbivores in groups. Groups were dominated by *C. sordidus*, which constituted ∼45% of all observed grouping individuals (figure 3) and had the highest percentage occurrence in groups (observed in ∼64% of all groups; figure 3).

Our assemblage-wide analysis of grouping behaviour among roving herbivores provided some support for the resource competition hypothesis. Territorial competitor densities had a positive effect on the grouping propensity of herbivores (β = 0.35, 95% CI = 0.12 to 0.60, logit scale; figure 2c; table S11) and a very weak effect on mean group sizes (β = 0.10, 95% CI = -0.00 to 0.21, log scale; figure 2d; table S12). The effect size of the latter relationship had a 97% posterior probability of being positive, but it was extremely small in magnitude.

In contrast, grouping behaviour showed strong positive density-dependence. Both grouping propensity (β = 0.77, 95% CI = 0.45 to 1.09, logit scale; figure 2e; table S11) and mean group sizes (β = 0.55, 95% CI = 0.42 to 0.67, log scale; figure 2f; table S12) of roving herbivores showed strong positive responses to increasing herbivore densities.

None of the results were qualitatively sensitive to the choice of priors and estimates of interest only varied by +-0.01 on using weakly informative priors. The results and inferences were robust to the exclusion of five repeat observations, removal of multi-species groups, and an interaction term between algae and structural complexity (ESM S5; tables S14-S28).

## 4. Discussion

Behavioural variation within and between populations can influence faunal-mediated ecosystem functions [3,12]. Incorporating this variation into current models and methods of assessing functional rates requires identifying causal links between behaviours and animals’ functional contributions [13], and identifying the mechanisms that drive this variation. Using a combination of individual-level observations and assemblage-scale surveys, we first demonstrated that variation in social behaviour can influence an individual’s functional contribution. We then identified drivers of variation in this social behaviour at both the individual and assemblage scale.

### (a) Individual behaviour-function covariation: Social foraging can facilitate herbivory

Our results demonstrate that the benefits of social foraging translate to higher per-capita herbivory – *C. sordidus* individuals in large groups (30 or more) had over 80% higher mean per-capita algal consumption potential than those feeding alone. We quantified this relationship only for *C. sordidus*, which was present in 64% of all observed groups during our censuses and made up for 45% of all grouping individuals (figure 3). While the magnitude of the relationship may vary by species, our preliminary observations of two other herbivorous fish species (*Scarus prasiognathus* and *Acanthurus auranticavus*) indicate that this may be a more general pattern across the herbivore assemblage *(personal observations)*. Thus, the same population of herbivores could differ substantially in their contribution to the function of herbivory depending on the distribution of grouping tendencies and group sizes in the population. More generally, this suggests that the expression of behavioural traits in a population can influence relevant ecosystem-scale functional rates. Several other studies, in both invertebrates and vertebrates, have demonstrated behaviour-function covariations [3,12,25]. Our results, together with these studies, highlight the importance of incorporating behaviour in assessments and predictions of faunal-mediated ecosystem functions, and explain why widely used static proxies, such as biomass, usually fail to reliably approximate functional rates [9,10].

While it has long been recognized that broad-scale patterns and processes are governed by processes operating at smaller scales [57], incorporating individual-level processes into large-scale assessments of ecosystem functioning is challenging in real-world conditions. Using social foraging by herbivorous coral reef fish as a model, our results demonstrate that investigations of the putative causes of behavioural variation can identify easily measurable drivers of this variation in functionally relevant behaviours. We believe this is essential for improving assessments of ecosystem functions and understanding how altered animal behaviours could affect ecosystem functions in rapidly changing environments.

### (b) Drivers of behavioural variation: Does resource competition or density-dependence drive herbivore grouping?

Our individual-scale analysis suggested that aggression from territorial competitors could be a cost to foraging for *C. sordidus* (figures 1e, 3). While the resource competition hypothesis predicts that non-territorial individuals overcome this cost by forming groups, our results provide only weak support for this prediction – increasing group sizes led to minimal reductions in aggression from territorial competitors – indicating that this mechanism may not be the primary mediator of the foraging benefit of grouping (figure 1f). This contrasts with previous studies which have attributed the foraging benefits of grouping to the dilution of aggression received from resource-defenders [32–34,58]. Invariably, most studies have considered resources defended by algae-farming damselfish – which strongly defend a highly productive resource [59]. However, on many reefs, algae-farming damselfish are not present in high densities and large areas of algal turfs, or the epilithic algal matrix, are defended by other less aggressive herbivores, such as surgeonfish [17,52,60]. Most of the aggressive encounters that *C. sordidus* individuals received in our observations were from these surgeonfish. However, the maximum aggression received, and variation in aggression, appeared to decrease in larger groups – potentially making aggression received in larger groups more predictable for participating individuals. This hypothesis can be tested by comparing the variability of aggression received in groups of different sizes. Animals modify their behaviour based on the predictability of resources and predation risk [61,62]. Similarly, they may benefit from higher predictability of potentially costly agonistic interactions while feeding. The adaptive benefits of social foraging are well documented across taxa, and the mechanisms through which they arise are variable [28]. An increase in feeding rates may result from mechanisms other than reduced aggression in larger groups, including a reduction in investment in vigilance [63], enhanced resource discovery [26,64], and generally from social information about resources and predation threat [65]. Further investigations to isolate the relative contributions of these mechanisms could be fruitful and could help us understand how environmental stressors influence the trade-offs involved in social foraging.

Scaling up to the entire assemblage of herbivores, our study provides some support for a reef-scale prediction of the resource-competition hypothesis – herbivores in areas with higher territorial competitor densities had higher grouping tendencies (figure 2c). However, mean group sizes only increased marginally (figure 2d). Together with our individual-level results – that the foraging benefit in larger groups was mediated only weakly by reduced aggression, these results suggest that interference competition, or competitive aggression, may not be as strong a driver of grouping at ecosystem scales as previously considered. Since the extent of competition depends on the degree of resource overlap [66], we expect grouping to be primarily driven by resource defence in situations where non-territorial individuals share a high degree of resource overlap with territorial individuals that strongly defend a limiting resource. In other cases, increasing group sizes beyond a certain point may provide diminishing benefits in accessing algal resources through dilution of aggression [67]. Previous experiments that removed territorial algae-farming damselfish to eliminate resource defence observed that the bite rates of solitary feeding roving herbivores in removal areas increased to a level comparable to those of schooling individuals [32,33]. While these elegant experiments provide strong evidence that grouping enables individuals to overcome resource defence and increase feeding rates inside territories, they do not explicitly test the hypothesis that resource competition causes group formation – which would predict a decrease in group formation on removal of territorial species.

Notably, our results suggest that the grouping behaviour of herbivorous coral reef fish is positively density-dependent, a finding consistent with evidence from laboratory and field studies of multiple taxa [68]. While evidence exists for density-dependence of group sizes [35], our findings indicate that grouping propensities – the likelihood of individuals joining groups – may also be positively density dependent. This suggests that social attraction and grouping behaviour can be constrained by density-dependent probabilistic processes (encounter rates between individuals) under natural conditions [68]. Similar density-dependence of grouping has been reported for pelagic fish, where rapid formation of large shoals occurs when a critical population density of loosely scattered individuals is achieved [69]. For animals which form both single-species and multi-species groups [23,30], at higher population densities, each individual may be more likely to encounter a partner or a group such that it can minimize the activity matching costs of grouping, while also accruing the benefits of group membership [36]. This might explain the increase in grouping propensities at high herbivore densities.

### (c) Behavioural variation and ecosystem functioning

Competitive and facilitative interactions are known to structure populations and communities in both terrestrial and marine systems, with flow on impacts on ecosystem functions [70]. If grouping tendencies vary in response to inter-specific competitive interactions between guilds, the spatial distribution of territoriality could have consequences for benthic community structure and functioning [71]. Since grouping appears to enhance herbivory, the spatial distribution of territoriality on a reef could lead to spatial heterogeneity of herbivory pressure [72], mediated by changes in grouping behaviour. However, these effects could vary based on the composition of the territorial fish assemblage, which would influence the degree of resource defence. A recent study found that the spatial variation in the aggression levels of territorial damselfish can influence fine-scale space use by other non-herbivorous fish [72], similar to the behaviour-mediated effects of predators on prey populations, which can lead to patchiness in herbivory [73,74]. Disturbance events, leading to loss of corals and structural complexity on reefs, can alter the abundance and aggressive behaviour of territorial fish [75]. This may further influence the rates and spatial distribution of herbivory, mediated by changes in grouping behaviour.

The positive density-dependence of grouping propensity and group sizes could have important consequences for functions associated with grouping, especially herbivory, making them particularly sensitive to the demography of herbivore populations. Since social foraging enhances per-capita herbivory, positive density-dependence of grouping indicates that individual foraging rates, and the function of herbivory, may decline non-linearly with depleting herbivore populations [25]. Similar effects have been previously described in coral reefs, where behavioural feedbacks in foraging behaviour lead to non-linear decreases in herbivory with rapid fishing because fish use social cues while foraging – which become less available at lower fish densities – giving rise to Allee effects [6,25]. In other words, at lower herbivore densities, fewer individuals might participate in groups and the resultant groups might be smaller. Thus, reduction in herbivory could be more severe than expected by most models using biomass as a proxy of herbivory – which would predict declines proportionate to biomass. Herbivores are significant determinants of producer community structure and productivity across systems – including savannas, salt marshes, and coral reefs [76–78]. In reefs subject to frequent coral mass mortalities, density-dependent reductions in herbivory could impede coral recovery by failing to control the growth of space occupying algae. [76].

C. sordidus was the most abundant in a large assemblage of interacting herbivores. Our results show that different functional groups of herbivores (scrapers, excavators, croppers) associate in multi-species social foraging networks (figure 3). These multi-functional associations have important implications for the spatial distribution of herbivory. For instance, herbivory on reefs may be restricted to relatively small patches, with different functional groups showing limited functional overlap, leading to high spatial complementarity [79]. However, herbivores forage across larger areas [17] and at higher feeding rates when in groups. This suggests that density-dependent social foraging could be important for expanding the spatial distribution and intensity of herbivory across different functional groups, thereby increasing functional redundancy. Studies on birds suggest that species associations in groups can change with structural complexity and resources [80] – both of which are rapidly changing on coral reefs [21]. Predicting how changes in group composition could influence functional roles requires understanding how the individual-level costs and benefits of grouping vary with group composition and the mechanisms governing group choice of animals – a promising avenue for future research [31,81]. Although extensive studies on mixed-species flocks of birds have generated insights on the causes of group formation, future work will benefit from also focusing on other taxa, including herbivorous fish [31] – given their functional relevance across marine ecosystems, and the opportunity to test the generality of our current understanding of group dynamics. While we tested the role of resource competition and density dependence on social foraging, future work will benefit from examining the effect of predation threat under natural settings since predation avoidance is a ubiquitous benefit of grouping [28]. For instance, coral reef fisheries often target large piscivorous fish, and how their extraction could influence functions is an important area of research [82].

Our analyses account for multiple confounding variables at the individual and transect-scale, and potential unobserved confounds at the site-scale, but there may be other unobserved confounds, particularly at the individual level. For example, both feeding behaviour and the tendency to participate in groups are influenced by the physiological state of an individual [65]. While such individual-level decisions may govern the mechanisms behind social foraging, they would not mask the observed relationships in our study.

## 5. Conclusion

Recent theoretical and empirical advancements have highlighted that animal behaviour can have profound impacts on ecosystem functioning [3,6,25,83]. Since behavioural modifications are often an animal’s first responses to anthropogenic stressors [84,85], understanding the links between animal behaviour and ecosystem function is essential to predict how ecosystem functions and services will respond to rapid environmental change. Our behavioural analyses quantify the individual-level benefits of grouping and how they translate to per-capita herbivory, and our assemblage-scale analyses identify easily measurable drivers of variation in grouping – linking individual behavioural variation with contributions to ecosystem functioning. This highlights the critical need to imagine ecosystems as dynamic networks of interacting, and behaving, organisms – whose actions contribute to the flow of energy. Incorporating this reality in quantitative frameworks has the potential to synthesize our understanding of organismal and functional ecology – for a more complete understanding of how our ecosystems function and how they will respond to rapid change.

## Acknowledgements

We thank NCBS – TIFR, and Nature Conservation Foundation for institutional, funding and administrative support. This work was supported by the Department of Atomic Energy, Government of India, Project Identification No. RTI 4006. We thank Fisheries Society of the British Isles for a Small Research Grant to HT (FSBI-RG23-555) which supported this work. We thank Dr. Sandeep Pulla and Dr. Jagdish Krishnaswamy for useful advice on the statistical analysis. Anwar Hussain, LakScuba Kavaratti, and ESMUC Kadmat provided immense logistical support for fieldwork. We are grateful to Mayukh Dey, Radhika Nair, and Herman Ramesh for discussions on this work. We acknowledge the support of the Department of Environment and Forests, Union Territory Lakshadweep for providing permission to carry out this study (F.No.1/5/2023-E&F/942). NCF received funding support from Shri AMM Murugappa Chettiar Research Centre (MCRC), Ashraya Hasta Trust, Cholamandalam Investment and Finance Company Ltd., Arvind Datar, and Muthukumaran Educational Trust. The Spanish National Research Council supported TA through the Memorandum of Understanding between Centre d’Estudis Avançats de Blanes (CEAB, CSIC) and Nature Conservation Foundation (NCF). RK was supported by a grant from National Geographic Society (NGS 96905R-22). Vectors for the graphics were used from Integration and Application Network, University of Maryland Media Library.

## Notes

### Competing Interest Statement

The authors have declared no competing interest.

