## Supplementary material for "Density-dependence and territorial competitors can modulate parrotfish social foraging and herbivory on coral reefs": SI

#### Supplementary Information

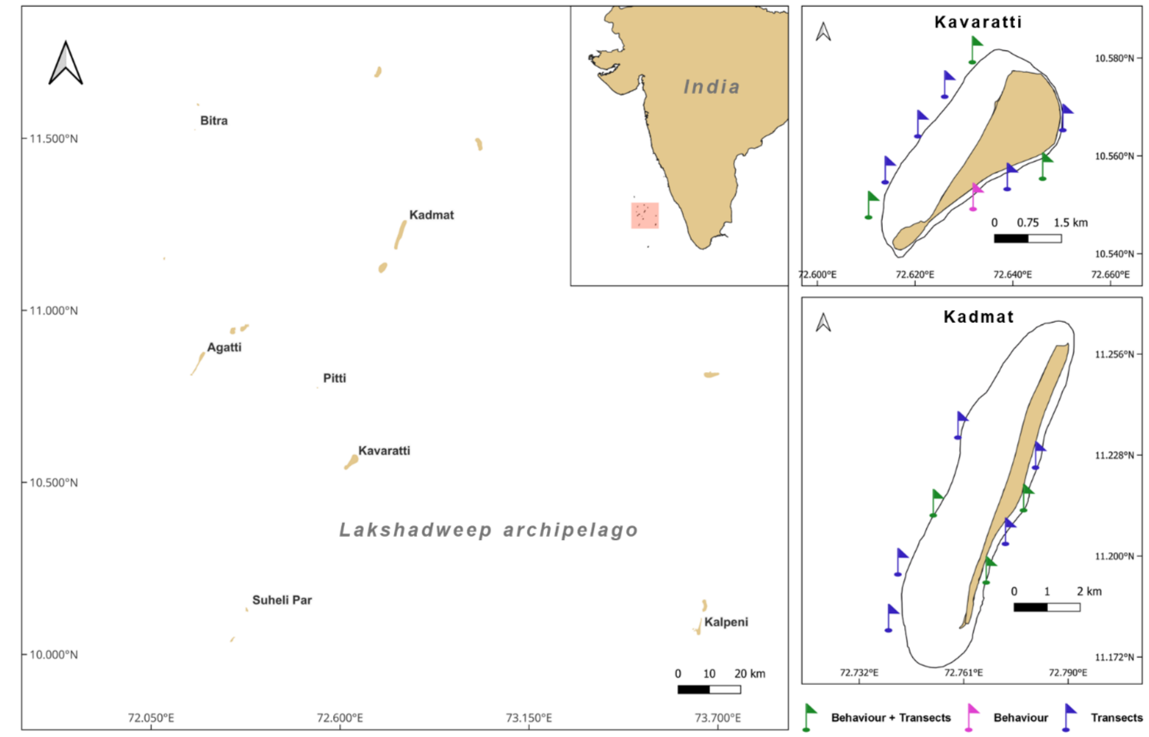


**Figure S1** Map of study region (left) and atolls sampled with sampling locations (right). At the locations marked in green, we conducted both – behavioural observations of *Chlorurus sordidus*, and community sampling; at the locations marked in pink, we only sampled behaviour; and at locations marked in blue, we only conducted community sampling. Black lines around the atolls indicate the boundary of the lagoons.

**Table S1:** Summary of scale of inference, objectives, sampling methods used to address the objectives, and the metrics of interest measured during sampling.

| Scale | Relationships of interest | Method (sample size) | Metrics of interest |
|---|---|---|---|
| Individual (*Chlorurus sordidus*) | • Effect of group size on individual feeding rates<br>• Effect of aggression received on individual feeding rates<br>• Effect of group size on aggression received from territorial competitors | Focal behavioural observations *of C. sordidus* in groups of different sizes (n = 103 focal individuals) | • Group sizes<br>• Bite rates<br>• Aggression rates received from territorial competitors |
| Herbivore assemblage | • Effect of territorial competitor densities on grouping behaviour of herbivores<br>• Effect of herbivore densities on grouping behaviour of herbivores | Underwater visual censuses using timed swims (n = 73 timed swims; 3 point counts in each swim) | • Herbivore densities – Number of roving herbivores per timed swim<br>• Territorial competitor densities – Number of territorial competitors per timed swim<br>• Grouping propensity – proportion of the herbivore assemblage observed in groups on a timed swim<br>• Mean group size of herbivores on a timed swim |

#### Section 1: Construction of DAGs

The process of constructing a DAG is independent of the collected data. Each directed edge in a DAG represents a hypothesised direct causal relationship between two variables, and the absence of an arrow conveys the assumption that the two variables are not directly causally related. We refer readers to the causal inference literature which provides more details on the utility and implementation of GCMs [1–11]. Since observational studies often try to address causal questions, DAGs help in transparently conveying causal assumptions about a system and make them open for scientific debate.

For the behavioural analyses, there are three relationships of interest represented in the DAG (Fig 1a of main text). All these relationships are at the scale of an individual *Chlorurus sordidus*. First, the direct effect of group size on bite rates (blue arrow); second, the direct effect of aggression on bite rates (red arrow); and third, the direct effect of group size on aggression rate (yellow arrow). Keeping these three relationships in mind, we identified variables that could affect any of these variables, and the pathways through which these effects could occur. This is represented in the form of directed arrows in the DAG. We did this based on a review of literature and our ecological knowledge. For instance, resource availability (algae) can affect the feeding rates (bite rates) of an individual and it can also influence the group sizes (Table S1). This makes resource availability, or algae, a confounder in the relationship between group size and bite rates. This is represented in DAG 1 by an arrow from the variable “Algae” pointing towards group size and another pointing towards bite rates. Similarly, the number of territorial individuals in the feeding area (Territorials) is a confounder when estimating the effect of group size on aggression rates. See Table S1 for the justification behind each arrow in DAG 1.

The variables we identified are potential confounders and mediators in the relationships of interest, operating at the scale of the focal individual or a group. However, our sampling design includes multiple sites, which may differ in other attributes – such as resource quality. Different sites may differ in nutrient content in algae because of differences in upwellings or pelagic nutrient inputs, leading to between-site differences in the quality of the algal resource. Resource quality at a site can influence group sizes, as well as bite rates – such that the optimal group size may increase and individuals may be able to feed at lower rates to meet nutritional requirements, allocating time to other activities. This means that there may be site-level unmeasured or unidentified confounders (such as differences in resource quality) – variables that affect both the exposure (group size) and outcome (bite rates). To account for this, we introduced site as a proxy variable for these potential confounders in the relationships of interest post-hoc [9] (see DAG 1 – arrows from “Site” to variables of interest). This assumes that these unobserved and unidentified confounding variables do not vary within a site but may vary between sites. There are multiple ways to deal with unobserved confounding in cross-sectional data [9]. We decided to use a Mundlak Device design [12], or a Group Mean Covariate design, because it lets us control for unmeasured site-level confounders, while also being able to account for site-level clustering in the data through random effects (random intercept). For this, site-level averages of a causal variable of interest are calculated and included as a predictor in the model – acting as a proxy for site-level unobserved confounders. Byrnes and Dee [9] provide a useful guide of how to deal with unobserved confounding variables in ecological studies, and we refer readers to their work for a detailed explanation.

We followed the procedure described above for constructing DAG 2 as well. There were four relationships of interest. First, the direct effect of herbivore densities on grouping propensity of roving herbivores; second, the direct effect of herbivore densities on mean group sizes of herbivores; third, the direct effect of territorial individual densities on grouping propensity; and fourth, the direct effect of territorial individual densities on mean group sizes. Here, the relationships of interest were at the scale of a timed swim. Like DAG 1, the variable "Site" in DAG 2 also represents potential unmeasured and unidentified site-level confounders between the relationships of interest. See Table S2 for the rationale behind each arrow in the DAG.

**Table S2**: Reasoning for all directed arrows in the directed acyclic graph (DAG) for *Chlorurus sordidus* foraging (Fig 1a).

| From | To | Justification |
|---|---|---|
| Aggression | Bite rate | Relationship of interest. Competitive interactions can negatively influence foraging [13,14]. |
| Algae | Bite rate | Resource availability can affect resource acquisition rates. |
| Algae | Aggression | Competitive aggression may be modulated by resource availability. Competition for resources may reduce if per capita resource availability is high. |
| Algae | Territorials | Higher resource levels may support higher aggressor abundance [15]. |
| Algae | Group size | Group sizes can be influenced by resource availability, or competition for resources. Group sizes can be constrained by competition for a limited resource [16] |
| Algae | Herbivores | Higher resources can support higher herbivore abundances [15]. |
| Territorials | Predators | Higher territorial fish abundance has the potential to support higher predator abundance. |
| Territorials | Aggression | Aggression rates received may be higher in areas where aggressor abundance is high (more potential aggressors). |
| Territorials | Group size | Number of territorial individuals in the vicinity may influence perceived aggression threat, and hence group sizes. |
| Group size | Bite rate | Relationship of interest. Animals in larger groups can have higher resource acquisition rates [17]. |
| Group size | Aggression | Relationship of interest. Per capita aggression received from competitors may be diluted in larger groups [13,14]. |
| Structure | Bite rate | Structure can influence bite rates by modulating perceived predation threat [18,19]. |
| Structure | Algae | Areas with more structural complexity could have more surface area for algae to grow. |
| Structure | Territorials | More structure means more surface area, more habitat, and potentially more refugia – which can influence abundances of site-attached territorial species. |

| | | |
|---|---|---|
| Structure | Group size | Structural complexity has been described to be typically associated with smaller group sizes, either due to visual hinderance or reduced perceived predation threat [20]. |
| Structure | Predators | Ambush predators rely on structural complexity – more structure may mean more predators[21]. |
| Herbivores | Group size | Group sizes may be dependent on the abundance or densities of potential participants [22,23]. |
| Herbivores | Predators | Higher herbivore abundance has the potential to support a higher predator abundance. |
| Predators | Bite rate | Predator abundance influences herbivory patterns and can influence bite rates by modulating perceived predation threat [18,19,24]. |
| Predators | Group size | Optimal group sizes may be sensitive to predation threat – larger groups may benefit through risk dilution or confusion effect, but may also be more detectable to predators [16,25,26] . |

**Table S3:** Reasoning for all directed arrows in the directed acyclic graph (DAG) for herbivore grouping (Fig 3a).

| **From** | **To** | **Justification** |
|---|---|---|
| Algae | Herbivores | Herbivore populations can respond to resource abundance [15]. |
| Algae | Territorials | Most territorial individuals in our study (*Acanthurus sp.*) are also herbivores, which track resource abundance [15]. |
| Algae | Grouping (Grouping propenstity/ Proportion grouping) | The costs and benefits of grouping can be dependent on resource abundance and distribution. Groups may be more efficient at finding resources, but suffer from the cost of increased competition – making grouping sensitive to resources [18]. |
| Algae | Mean group size | Higher resources may support larger groups without the costs of intra-group competition outweighing the benefits. |
| Herbivores | Predators | Higher prey abundance can support higher predator abundance. |
| Herbivores | Mean group size | Relationship of interest, group sizes can be density dependent [22,23]. |
| Herbivores | Grouping (Grouping propensity/ Proportion grouping) | Relationship of interest. Grouping behaviour can be density dependent [27]. |
| Structure | Algae | More structural complexity can provide more area for algae to grow. |
| Structure | Predators | Several ambush predators require high structural complexity [21]. |
| Structure | Mean group size | Since grouping behaviour provides predation avoidance benefits [28], it can be sensitive to perceived predation threat. Higher structural complexity can reduce perceived predation threat since it provides more refuges [18,19]. |
| Structure | Grouping (Grouping propenstity/ Proportion grouping) | Since grouping behaviour provides predation avoidance benefits [28], it can be sensitive to perceived predation threat. Higher structural complexity can reduced perceived predation threat since it provides more refuges [18,19]. |
| Structure | Territorials | Higher structural complexity can support higher herbivore abundances by increasing surface area for algal growth and providing more refugia [15]. Most territorial individuals in our study were herbivores. |
| Territorials | Predators | Territorial individuals can be potential prey for predators, which may track prey abundance. |
| Territorials | Mean group size | Relationship of interest. Larger groups may be more efficient at accessing defended resources [14,29]. |

| | | |
|---|---|---|
| Territorials | Grouping (Grouping propenstity/ Proportion grouping) | Relationship of interest. The resource competition hypothesis [13,14] predicts that a larger proportion of the herbivore population should participate in areas of higher resource defence. |
| Predators | Mean group size | Grouping behaviour can change in response to perceived predation threat [26]. |
| Predators | Grouping (Grouping propenstity/ Proportion grouping) | Grouping behaviour can change in response to perceived predation threat [26]. |

#### Section 2: Testing conditional independences

We validated our DAGs by testing the implied conditional independencies in our observational dataset using the R package *“dagitty”* [30].

Most DAGs imply a set of conditional independences – assuming the relationships specified in the DAG, there can be a possible set of statements describing which observed variables should be independent of one another – both with and without conditioning on other variables [3,4]. These implications are testable in the data and are useful in testing the assumptions of the DAG. For example, one of the implications of DAG 1 is *Bite rate _||_ Territorials | Algae , Aggression, Predators, Group size, Structure.* This means that, according to our DAG, bite rate should be independent of total territorial individuals after conditioning on algae, aggression, predators, group size, and structure. This effectively translates to saying that after controlling for these variables, there is no path for territorials and bite rates to be related. This implication is testable because data on all these variables is available. The package “dagitty” has functions to perform the entire operation – from identifying the implied conditional independences to testing them. In this example, the functions would condition on the set of variables and determine the residual correlation between bite rate and territorials. If there is no remaining correlation, it would mean that the data supports the assumptions about the relationships between bite rates and total territorial individuals specified in our DAG. Conversely, if there is a strong residual correlation after conditioning, it would mean that there might be other pathways for bite rate and territorial abundance to be related besides those specified in the DAG, and would indicate that the data is not consistent with the causal assumptions depicted in the DAG.

In the absence of standardised methods for testing conditional independences between continuous and unordered categorical variables [31], we tested conditional independences implied by the DAG without the site variable.

The code accompanying the manuscript provides annotated steps on performing these tests for both the DAGs. The results of these tests are presented in Table S3 and Table S4. There is no strong residual correlation after conditioning on the relevant variables – suggesting that our data and DAGs are compatible, or, in other words, the data supports the conditional independences implied by the DAGs.

**Table S4:** Conditional independences tested for DAG 1 representing individual level *Chlorurus sordidus* behaviour

| Conditional independence | Correlation coefficient after conditioning | 95% Confidence Interval |
|---|---|---|
| Bite rate _||_ Territorials \| Algae , Aggression, Predators, Group size, Structure | 0.06 | -1.34, 0.26 |
| Aggression _||_ Predators \| Algae, Group size, Territorials | -0.13 | -0.32,0.06 |
| Aggression _||_ Structure \| Algae, Group size, Territorials | 0.22 | 0.03, 0.40 |

**Table S5:** Conditional independences tested for DAG 2 representing grouping in the roving herbivore assemblage

| Conditional independence | Correlation coefficient after conditioning | 95% Confidence Interval |
|---|---|---|
| Algae _||_ Predators \| Herbivores, Structure, Territorials | 0.20 | -0.03, 0.41 |
| Mean group size _||_ Proportion Grouping \| Algae, Predators, Herbivores, Structure, Territorials | -0.02 | -0.22,0.25 |
| Herbivores _||_ Structure \| Algae | -0.27 | -0.47, -0.04 |
| Herbivores _||_ Territorials \| Algae | -0.05 | -0.28, 0.18 |

#### Section 3: Deriving statistical models from DAGs

DAGs help in avoiding statistical biases (like collider, confounding, overcontrol) which may lead to erroneous estimates when using model selection methods meant for prediction, or causal salad models that include all variables hypothesised to influence the response variable in a statistical model [32].

We used the web tool "*dagitty*" *(www.dagitty.net)[30]* to derive statistical models, or a set of adjustment variables required to estimate effects of interest, from our DAGs. *dagitty* automates the process of identifying variables that need to be incorporated into a statistical model to block biasing paths. However, this process can also be performed manually by selecting variables such that all paths between the two variables, other than the direct path of interest, are blocked. An easy explanation can be found in Pearl [4] and on "https://www.dagitty.net/learn/". We detail this process for our models below.

##### *(a) Model 1*

$$Bite\ rate_i \sim Normal\ (\mu_i, \sigma)$$

$$\mu_i = \alpha_{site} + \beta_1 Group\ size + \beta_2 Aggression\ received + \beta_3 Algae + \beta_4 Structure + \beta_5 \overline{Group\ size}_{Site}$$

To estimate the direct effect of group sizes on bite rates, our model needed to adjust for four confounders shown in the DAG – algae, structure, predators, and site – and one mediator – aggression received. Blocking the mediator is required when estimating a direct effect, but not a total effect.

The group mean covariate design controls for the "site" variable by using the site-level average of group sizes in the model. It is important to note that our measure or "Predators" for the focal follows of *C. sordidus* was at the site-scale. This effectively creates an arrow from "Site" to "Predators" in our DAG. Thus, including both, the site level average for group size and predators, in the model would open biasing paths, or is statistically redundant. Hence, we only used the site-level average of group sizes in our model and excluded predators. Additionally, predator abundance was measured using separate 50 m x 10 m belt transects and not on the focal follows.

This model was also compatible with estimating the direct effect of aggression on bite rates. In this relationship, the confounders are group size, algae, structure, and site. While there is no direct arrow from structure to aggression in the DAG, it is a confounder through the path Structure -> Territorials -> Aggression. This conveys the assumption that while structure does not directly influence aggression, it influences the abundance of territorial individuals which influence aggression rates. Hence, to block this path, we could have included either, structure or territorial abundance, in the model.

##### *(b) Model 2*

$$Aggression\ rate\ received \sim HurdleGamma(hu, mu, scale)$$

$$logit(hu) = \alpha_{site\_h} + \beta_1 Group\ size + \beta_2 Aggressors + \beta_3 Algae + \beta_4 \overline{Group\ size}_{Site}$$

$$\log(mu) = \alpha_{site\_m} + \beta_5 Group\ size + \beta_6 Aggressors + \beta_7 Algae + \beta_8 \overline{Group\ size}_{Site}$$

To estimate the direct effect of group size on aggression rate received, our model needed to adjust for the confounders group size, aggressor abundance, algae, and group size – variables which influence both aggression received and group size.

##### *(c) Model 3 and Model 4*

$$Grouping\ propensity_i \sim Beta\ (\mu_i, \varphi)$$

$$logit(\mu_i) = \alpha_{site} + \beta_1 Herbivore\ density + \beta_2 Territorial\ competitor\ density + \beta_3 Algae + \beta_4 Structure + \beta_5 Predators + \beta_6 \overline{Herbivore\ density}_{Site}$$

$$Mean\ group\ size_i \sim Gamma\ (\mu_i, shape)$$

$$log(\mu_i) = \alpha_{site} + \beta_1 Herbivore\ density + \beta_2 Territorial\ competitor\ density + \beta_3 Algae + \beta_4 Structure + \beta_5 Predators + \beta_6 \overline{Herbivore\ density}_{Site}$$

These models block a mediator (number of predators) and other biasing paths through algae and structure.

#### Section 4: Selection of priors

We used informative priors for all parameters where variables were expected to have a direct effect on the outcome variable in our DAG. However, since we do not borrow our priors from the posterior estimates of similar studies in related taxa (which, to our knowledge, are not available for many parameters), we did not use very narrow priors. The priors we used indicate expected directions of the effects of interest but leave enough probability on the opposite side of zero for the priors to not restrict the posterior to either positive or negative values. The priors for the site level averages, which adjust for potential site-level confounders, were set to Normal(0,10).

***(a) Model 1:***

$$Bite\ rate_i \sim Normal\ (\mu_i, \sigma)$$

$$\mu_i = \alpha_{site} + \beta_1 Group\ size + \beta_2 Aggression\ received + \beta_3 Algae + \beta_4 Structure + \beta_5 \overline{Group\ size}_{Site}$$

$$\beta_{1,3} \sim Normal\ (1,1)$$

$$\beta_2 \sim Normal\ (-1,1)$$

$$\beta_4 \sim Normal\ (0,1)$$

$$\alpha_{site} \sim Half - Normal\ (\bar{\alpha}_{site}, \sigma_{site})$$

**Table S6:** Justification for priors used in model 1.

| Parameter | Prior | Justification |
|---|---|---|
| $\beta_1$ | $Normal\ (1,1)$ | Group size can increase resource acquisition rates [17] |
| $\beta_2$ | $Normal\ (-1,1)$ | Receiving aggression can be a foraging cost. Animals face trade-offs in time investment in different behaviours [33]. |
| $\beta_3$ | $Normal\ (1,1)$ | Higher resources can lead to higher resource acquisition rates since time required to find resources is reduced (if resource was patchy), or it may also increase if resource was limiting . |
| $\beta_4$ | $Normal\ (0,1)$ | Structural complexity can modify perceived predation threat, leading to similar effects [18]. |

***(b) Model 2:***

$$Aggression\ rate\ received \sim HurdleGamma(hu, mu, scale)$$

$$logit(hu) = \alpha_{site\_h} + \beta_1 Group\ size + \beta_2 Aggressors + \beta_3 Algae + \beta_4 \overline{Group\ size}_{Site}$$

$$\log(mu) = \alpha_{site\_m} + \beta_5 Group\ size + \beta_6 Aggressors + \beta_7 Algae + \beta_8 \overline{Group\ size}_{Site}$$

$$\beta_{1,3,6} \sim Normal\ (1,1)$$

$$\beta_{2,5,7} \sim Normal\ (-1,1)$$

$$\alpha_{site_{h,m}} \sim Half - Normal\left(\bar{\alpha}_{site_{h,m}}, \sigma_{site_{h,m}}\right)$$

**Table S7:** Justification for priors used in model 2.

| Parameter | Prior | Justification |
|---|---|---|
| $\beta_1$ | $Normal\ (1,1)$ | Dilution of aggression in larger groups [14]. |
| $\beta_2$ | $Normal\ (-1,1)$ | When feeding in areas where potential aggressors/ territorial competitors are more abundant, the chances of receiving aggression may be higher. |
| $\beta_3$ | $Normal\ (1,1)$ | When resources are more abundant, competition may be relaxed – possibly leading to lower aggression. |
| $\beta_5$ | $Normal\ (-1,1)$ | Dilution of aggression in larger groups [14]. |
| $\beta_6$ | $Normal\ (1,1)$ | When feeding in areas where potential aggressors/ territorial competitors are more abundant, the chances of receiving aggression may be higher. |
| $\beta_7$ | $Normal\ (-1,1)$ | When resources are more abundant, competition may be relaxed – possibly leading to lower aggression. |

##### *(c) Model 3:*

$$Grouping\ propensity_i \sim Beta\ (\mu_i, \varphi)$$

$$logit(\mu_i) = \alpha_{site} + \beta_1 Herbivore\ density + \beta_2 Territorial\ competitor\ density + \beta_3 Algae + \beta_4 Structure + \beta_5 Predators + \beta_6 \overline{Herbivore\ density}_{Site}$$

$$\beta_{1,2,5} \sim Normal\ (1,1)$$

$$\beta_3 \sim Normal\ (0,1)$$

$$\beta_4 \sim Normal\ (-1,1)$$

$$\alpha_{site} \sim Normal\ (\bar{\alpha}_{site}, \sigma_{site})$$

**Table S8:** Justification for priors used in model 3.

| Parameter | Prior | Justification |
|---|---|---|
| $\beta_1$ | $Normal\ (1,1)$ | At higher herbivore densities, chances of encountering suitable group partners can be higher [22,34] |
| $\beta_2$ | $Normal\ (1,1)$ | When resource defence is higher (more resource defenders/ territorial competitors), grouping propensities may be higher to gain access to defended resources – a prediction of the resource competition hypothesis [13,14]. |
| $\beta_3$ | $Normal\ (0,1)$ | If grouping is constrained by resources (cost of intra-group competition is high while grouping), increasing resources can lead to increase in grouping propensities. Conversely, if animals were primarily grouping for enhanced resource discovery in groups, increased resources (or lower patchiness) may reduce grouping propensities. |
| $\beta_4$ | $Normal\ (-1,1)$ | Perceived predation threat can be lower in areas of higher structural complexity; and visual hinderance may limit grouping propensities. |
| $\beta_5$ | $Normal\ (1,1)$ | Grouping is well established to be provide anti-predator benefits, and grouping propensities may be higher in areas of higher perceived predation threat (predator abundance) [25]. |

##### *(d) Model 4:*

$$Mean\ group\ size_i \sim Gamma\ (\mu_i, shape)$$

$$log(\mu_i) = \alpha_{site} + \beta_1 Herbivore\ density + \beta_2 Territorial\ competitor\ density + \beta_3 Algae + \beta_4 Structure + \beta_5 Predators + \beta_6 \overline{Herbivore\ density}_{Site}$$

$$\beta_{1,2,5} \sim Normal\ (1,1)$$

$$\beta_3 \sim Normal\ (0,1)$$

$$\beta_4 \sim Normal\ (-1,1)$$

$$\alpha_{site} \sim Normal\ (\bar{\alpha}_{site}, \sigma_{site})$$

**Table S9:** Justification for priors used in model 4.

| Parameter | Prior | Justification |
|---|---|---|
| $\beta_1$ | $Normal\ (1,1)$ | At higher herbivore densities, there are higher chances of encountering more suitable group partners such that activity matching costs can be minimized while also accruing the benefits of larger groups [22,34]. |
| $\beta_2$ | $Normal\ (1,1)$ | When resource defence is higher (more resource defenders/ territorial competitors), group sizes may be higher to gain access to defended resources. |
| $\beta_3$ | $Normal\ (0,1)$ | If group sizes are constrained by resources (cost of competition is high while grouping), increasing resources can lead to increase in group sizes. Conversely, if animals were primarily grouping for enhanced resource discovery in groups, higher resources (or lower patchiness), may reduce group sizes. |
| $\beta_4$ | $Normal\ (-1,1)$ | Perceived predation threat can be lower in areas of higher structural complexity, therefore reducing group sizes; and visual hinderance may limit group sizes in more complex habitats. |
| $\beta_5$ | $Normal\ (1,1)$ | Grouping is well established to be provide anti-predator benefits, and group sizes may be higher in areas of higher perceived predation threat (predator abundance) [25]. |

### Model output

**Table S10:** Model output for model 1 of main text (bite rates of *C. sordidus*)

| **Multilevel hyperparameters: Site (number of levels: 6)** | | | |
|---|---|---|---|
| **Parameter** | **Estimate** | **Lower 95% CI** | **Upper 95% CI** |
| sd(Intercept) | 0.40 | 0.03 | 1.26 |
| **Regression Coefficients** | | | |
| Intercept | 0.00 | -0.44 | 0.45 |
| Group size | 0.50 | 0.33 | 0.66 |
| Aggression rate | -0.29 | -0.46 | -0.12 |
| Algae | 0.14 | -0.08 | 0.36 |
| Structure | -0.07 | -0.28 | 0.13 |
| Mean group size at site | -0.25 | -0.72 | 0.21 |
| All coefficient estimates are for scaled outcome (bite rate) and predictor variables. | | | |

**Table S11:** Model output for model 2 of main text (aggression rates)

| **Multilevel hyperparameters: Site (number of levels: 6)** | | | |
|---|---|---|---|
| **Parameter** | **Estimate** | **Lower 95% CI** | **Upper 95% CI** |
| sd(Intercept) | 0.26 | 0.01 | 0.95 |
| sd(Hurdle intercept) | 0.82 | 0.05 | 2.34 |
| **Regression Coefficients : Hurdle** | | | |
| Intercept | -0.52 | -1.38 | 0.39 |
| Group size | 0.29 | -0.15 | 0.75 |
| Algae | -0.34 | -0.90 | 0.26 |
| Total aggressors | -0.16 | -0.64 | 0.32 |
| Mean group size at site | -0.05 | -1.05 | 0.90 |
| **Regression Coefficients: Gamma** | | | |
| Intercept | -0.28 | -0.63 | 0.04 |
| Group size | -0.15 | -0.30 | 0.01 |
| Algae | 0.06 | -0.15 | 0.28 |
| Total aggressors | 0.14 | -0.05 | 0.33 |
| Mean group size at site | -0.02 | -0.30 | 0.38 |
| All coefficients estimates are for scaled predictor variables. The coefficients for the hurdle model are on the logit scale, and coefficients for the gamma part are on the log scale | | | |

**Table S12:** Model output for model 3 of main text (proportion grouping)

| **Multilevel hyperparameters: Site (number of levels: 15)** | | | |
|---|---|---|---|
| **Parameter** | **Estimate** | **Lower 95% CI** | **Upper 95% CI** |
| sd(Intercept) | 0.23 | 0.01 | 0.59 |
| **Regression Coefficients** | | | |
| Intercept | 1.16 | 0.91 | 1.41 |
| Herbivores | 0.77 | 0.45 | 1.09 |
| Territorials | 0.35 | 0.12 | 0.60 |
| Algae | 0.05 | -0.20 | 0.28 |
| Structure | -0.20 | -0.42 | 0.02 |
| Predators | 0.02 | -0.23 | 0.32 |
| Mean herbivore density at site | -0.06 | -0.35 | 0.23 |
| All coefficient estimates are on the logit scale and correspond to scaled predictor variables. | | | |

**Table S13:** Model output for model 4 of main text (mean group sizes)

| **Multilevel hyperparameters: Site (number of levels: 15)** | | | |
|---|---|---|---|
| **Parameter** | **Estimate** | **Lower 95% CI** | **Upper 95% CI** |
| sd(Intercept) | 0.09 | 0.00 | 0.25 |
| **Regression Coefficients** | | | |
| Intercept | 2.05 | 1.94 | 2.16 |
| Herbivores | 0.55 | 0.42 | 0.67 |
| Territorials | 0.10 | -0.00 | 0.21 |
| Algae | 0.14 | 0.03 | 0.24 |
| Structure | -0.08 | -0.19 | 0.03 |
| Predators | -0.08 | -0.19 | 0.03 |
| Mean herbivore density at site | 0.17 | 0.03 | 0.31 |
| All coefficient estimates are on the log scale and correspond to scaled predictor variables. | | | |

#### Section 5: Sensitivity analyses

We performed a series of sensitivity analyses to test the robustness of our inferences to certain assumptions and exclusion of some data. First, in the focal behavioural observations, if an individual moved to a different group, it was recorded as a separate observation (n = 5). We removed these repeat observations and refit all models. Second, we removed multi-species groups from our behavioural analyses because the benefits of grouping may vary with group composition [35]. Third, resource (EAM cover) was quantified as a planar percentage but the available area for algae to grow can vary with structural complexity. Thus, we repeated all models with an interaction between algal cover and structural complexity – allowing the effect of resource to change with structural complexity. Fourth, we tested the sensitivity of our inferences to the choice of priors by refitting all models with the priors for coefficients set to Normal(0,10) and default priors for all other parameters.

##### (a) Sensitivity Analysis 1: Removing repeat observations from behavioural analyses

**Table S14:** Model 1 results for sensitivity analysis 1

| **Multilevel hyperparameters: Site (number of levels: 6)** | | | |
|---|---|---|---|
| **Parameter** | **Estimate** | **Lower 95% CI** | **Upper 95% CI** |
| sd(Intercept) | 0.41 | 0.03 | 1.29 |
| **Regression Coefficients** | | | |
| Intercept | 0.00 | -0.46 | 0.45 |
| Group size | 0.49 | 0.32 | 0.66 |
| Aggression rate | -0.29 | -0.46 | -0.12 |
| Algae | 0.16 | -0.11 | 0.37 |
| Structure | -0.13 | -0.33 | 0.08 |
| Mean group size at site | -0.21 | -0.68 | 0.23 |
| All coefficient estimates are for scaled outcome (bite rate) and predictor variables. | | | |

**Table S15:** Model 2 results for sensitivity analysis 1

| Multilevel hyperparameters: Site (number of levels: 6) | | | |
|---|---|---|---|
| **Parameter** | **Estimate** | **Lower 95% CI** | **Upper 95% CI** |
| sd(Intercept) | 0.26 | 0.01 | 0.92 |
| sd(Hurdle intercept) | 1.01 | 0.08 | 2.75 |
| **Regression Coefficients : Hurdle** | | | |
| Intercept | -0.64 | -1.64 | 0.40 |
| Group size | 0.38 | -0.09 | 0.88 |
| Algae | -0.37 | -0.97 | 0.28 |
| Total aggressors | -0.01 | -0.51 | 0.50 |
| Mean group size at site | -0.01 | -1.17 | 1.16 |
| **Regression Coefficients: Gamma** | | | |
| Intercept | -0.29 | -0.65 | 0.04 |
| Group size | -0.15 | -0.31 | 0.01 |
| Algae | 0.06 | -0.16 | 0.28 |
| Total aggressors | 0.15 | -0.05 | 0.35 |
| Mean group size at site | 0.02 | -0.30 | 0.38 |
| All coefficients estimates are for scaled predictor variables. The coefficients for the hurdle model are on the logit scale, and coefficients for the gamma part are on the log scale | | | |

**Sensitivity Analysis 2: Removing multi-species groups from behavioural analyses**

Our focal behavioural observations of *Chlorurus sordidus* consisted of individuals in both mono-specific (n=65) and multi-species (n = 27) groups. Here, we considered a group which had at least one individual of any species other than *C. sordidus* to be a multi-species group. However, there can be considerable variation in group composition between multi-species groups. For instance, a group of 15 individuals may have 14 *C. sordidus* and 1 *Scarus psittacus*, or 9 C. sordidus and 6 individuals of other species. Both can be considered multi-specific, but the group context experienced by the focal individual would be different. Notably, the benefits of grouping may vary with group composition [40]. This sensitivity analysis aimed to test the sensitivity of our inferences to changes in benefits accrued by an individual depending on group composition.

Specifically, it addresses whether our results are robust to the exclusion of multi-species groups. In our data, multi-species groups of *C. sordidus* were dominated by C. soridus itself – hence it was difficult to establish a Boolean condition of what qualifies as a mono-specific or multi-specific group. Thus, we used proportion of individuals belonging to *C. sordidus* as a proxy for "mono-specificity" of a group. We created three arbitrary thresholds of what qualifies as a mono-specific group depending on the proportion of individuals in a group of the species *C. sordidus* and repeated all behavioural analysis with only those observations. We sequentially considered groups to be mono-specific if the proportion of individuals that were *C. soridus* were over 50%, 80%, and 95% in groups. This resulted in 101, 87, and 74 remaining observations respectively.

Tables S16-S21 show the model outputs. There was some increase in the coefficient that represents the effect of group size on bite rates, but this does not change any inferences drawn from our results.

**Table S16:** Results from model 1 (bite rate) with only mono-specific groups at a 50% threshold

| **Multilevel hyperparameters: Site (number of levels: 6)** | | | |
|---|---|---|---|
| **Parameter** | **Estimate** | **Lower 95% CI** | **Upper 95% CI** |
| sd(Intercept) | 0.43 | 0.03 | 1.34 |
| **Regression Coefficients** | | | |
| Intercept | 0.00 | -0.48 | 0.49 |
| Group size | 0.50 | 0.32 | 0.67 |
| Aggression rate | -0.29 | -0.46 | -0.11 |
| Algae | 0.14 | -0.09 | 0.35 |
| Structure | -0.07 | -0.28 | 0.13 |
| Mean group size at site | -0.24 | -0.73 | 0.23 |
| All coefficient estimates are for scaled outcome (bite rate) and predictor variables. | | | |

**Table S17:** Results from model 2 (aggression rate) with only mono-specific groups at a 50% threshold

| **Multilevel hyperparameters: Site (number of levels: 6)** | | | |
|---|---|---|---|
| **Parameter** | **Estimate** | **Lower 95% CI** | **Upper 95% CI** |
| sd(Intercept) | 0.26 | 0.01 | 0.95 |
| sd(Hurdle intercept) | 0.97 | 0.09 | 2.56 |
| **Regression Coefficients: Hurdle** | | | |
| Intercept | -0.59 | -1.57 | 0.41 |
| Group size | 0.33 | -0.13 | 0.80 |
| Algae | -0.39 | -0.98 | 0.22 |
| Total aggressors | -0.09 | -0.58 | 0.39 |
| Mean group size at site | 0.16 | -0.68 | 1.18 |
| **Regression Coefficients: Gamma** | | | |
| Intercept | -0.29 | -0.64 | 0.05 |
| Group size | -0.15 | -0.30 | 0.02 |
| Algae | 0.07 | -0.15 | 0.28 |
| Total aggressors | 0.14 | -0.05 | 0.34 |
| Mean group size at site | -0.01 | -0.36 | 0.31 |
| All coefficients estimates are for scaled predictor variables. The coefficients for the hurdle model are on the logit scale, and coefficients for the gamma part are on the log scale | | | |

**Table S18:** Results from model 1 (bite rate) with only mono-specific groups at a 80% threshold

| **Multilevel hyperparameters: Site (number of levels: 6)** | | | |
|---|---|---|---|
| **Parameter** | **Estimate** | **Lower 95% CI** | **Upper 95% CI** |
| sd(Intercept) | 0.36 | 0.02 | 1.17 |
| **Regression Coefficients** | | | |
| Intercept | 0.01 | -0.39 | 0.43 |
| Group size | 0.54 | 0.37 | 0.72 |
| Aggression rate | -0.35 | -0.53 | -0.16 |
| Algae | 0.19 | -0.05 | 0.42 |
| Structure | -0.04 | -0.26 | 0.17 |
| Mean group size at site | -0.26 | -0.70 | 0.16 |
| All coefficient estimates are for scaled outcome (bite rate) and predictor variables. | | | |

**Table S19:** Results from model 2 (aggression rate) with only mono-specific groups at a 80% threshold

| **Multilevel hyperparameters: Site (number of levels: 6)** | | | |
|---|---|---|---|
| **Parameter** | **Estimate** | **Lower 95% CI** | **Upper 95% CI** |
| sd(Intercept) | 0.35 | 0.02 | 1.16 |
| sd(Hurdle intercept) | 1.03 | 0.09 | 2.77 |
| **Regression Coefficients: Hurdle** | | | |
| Intercept | -0.60 | -1.61 | 0.47 |
| Group size | 0.38 | -0.11 | 0.92 |
| Algae | -0.40 | -1.05 | 0.28 |
| Total aggressors | -0.09 | -0.64 | 0.44 |
| Mean group size at site | 0.06 | -1.29 | 1.15 |
| **Regression Coefficients: Gamma** | | | |
| Intercept | -0.28 | -0.71 | 0.13 |
| Group size | -0.12 | -0.29 | 0.06 |
| Algae | 0.04 | -0.21 | 0.29 |
| Total aggressors | 0.14 | -0.07 | 0.36 |
| Mean group size at site | 0.08 | -0.31 | 0.52 |
| All coefficients estimates are for scaled predictor variables. The coefficients for the hurdle model are on the logit scale, and coefficients for the gamma part are on the log scale | | | |

**Table S20:** Results from model 1 (bite rate) with only mono-specific groups at a 95% threshold

| **Multilevel hyperparameters: Site (number of levels: 6)** | | | |
|---|---|---|---|
| **Parameter** | **Estimate** | **Lower 95% CI** | **Upper 95% CI** |
| sd(Intercept) | 0.50 | 0.04 | 1.52 |
| **Regression Coefficients** | | | |
| Intercept | 0.03 | -0.50 | 0.59 |
| Group size | 0.53 | 0.32 | 0.74 |
| Aggression rate | -0.35 | -0.55 | -0.15 |
| Algae | 0.16 | -0.10 | 0.41 |
| Structure | 0.03 | -0.22 | 0.28 |
| Mean group size at site | -0.25 | -0.82 | 0.28 |
| All coefficient estimates are for scaled outcome (bite rate) and predictor variables. | | | |

**Table S21:** Results from model 2 (aggression rate) with only mono-specific groups at a 95% threshold

| **Multilevel hyperparameters: Site (number of levels: 6)** | | | |
|---|---|---|---|
| **Parameter** | **Estimate** | **Lower 95% CI** | **Upper 95% CI** |
| sd(Intercept) | 0.39 | 0.02 | 1.27 |
| sd(Hurdle intercept) | 1.83 | 0.50 | 4.09 |
| **Regression Coefficients: Hurdle** | | | |
| Intercept | -0.68 | -2.12 | 0.90 |
| Group size | 1.18 | 0.48 | 2.00 |
| Algae | -0.37 | -1.17 | 0.43 |
| Total aggressors | -0.32 | -1.01 | 0.34 |
| Mean group size at site | -0.23 | -2.21 | 1.62 |
| **Regression Coefficients: Gamma** | | | |
| Intercept | -0.22 | -0.69 | 0.24 |
| Group size | -0.21 | -0.50 | 0.08 |
| Algae | -0.04 | -0.31 | 0.23 |
| Total aggressors | 0.16 | -0.08 | 0.40 |
| Mean group size at site | 0.07 | -0.40 | 0.58 |
| All coefficients estimates are for scaled predictor variables. The coefficients for the hurdle model are on the logit scale, and coefficients for the gamma part are on the log scale | | | |

**Sensitivity Analysis 3: Interaction between algae and structural complexity**

**Table S22:** Results from model 1 (bite rate) with an interaction between algae and structural complexity

| **Multilevel hyperparameters: Site (number of levels: 6)** | | | |
|---|---|---|---|
| **Parameter** | **Estimate** | **Lower 95% CI** | **Upper 95% CI** |
| sd(Intercept) | 0.43 | 0.03 | 1.35 |
| **Regression Coefficients** | | | |
| Intercept | -0.01 | -0.47 | 0.47 |
| Group size | 0.50 | 0.33 | 0.67 |
| Aggression rate | -0.30 | -0.47 | -0.12 |
| Algae | 0.14 | -0.09 | 0.36 |
| Structure | -0.07 | -0.28 | 0.13 |
| Mean group size at site | -0.25 | -0.73 | 0.21 |
| Algae: Structure | 0.03 | -0.16 | 0.21 |
| All coefficient estimates are for scaled outcome (bite rate) and predictor variables. | | | |

**Table S23:** Model output for model 3 (proportion grouping) with an interaction between algae and structural complexity

| **Multilevel hyperparameters: Site (number of levels: 15)** | | | |
|---|---|---|---|
| **Parameter** | **Estimate** | **Lower 95% CI** | **Upper 95% CI** |
| sd(Intercept) | 0.24 | 0.01 | 0.62 |
| **Regression Coefficients** | | | |
| Intercept | 1.16 | 0.90 | 1.42 |
| Herbivores | 0.77 | 0.46 | 1.10 |
| Territorials | 0.35 | 0.12 | 0.59 |
| Algae | 0.06 | -0.20 | 0.29 |
| Structure | -0.20 | -0.41 | 0.02 |
| Predators | 0.03 | -0.22 | 0.32 |
| Mean herbivore density at site | -0.08 | -0.38 | 0.22 |
| Algae: Structure | 0.06 | -0.15 | 0.28 |
| All coefficient estimates are on the logit scale and correspond to scaled predictor variables. | | | |

**Table S24:** Model output for model 4 (mean group sizes) with an interaction between algae and structural complexity

| **Multilevel hyperparameters: Site (number of levels: 15)** | | | |
|---|---|---|---|
| **Parameter** | **Estimate** | **Lower 95% CI** | **Upper 95% CI** |
| sd(Intercept) | 0.08 | 0.00 | 0.23 |
| **Regression Coefficients** | | | |
| Intercept | 2.04 | 1.94 | 2.15 |
| Herbivores | 0.55 | 0.43 | 0.67 |
| Territorials | 0.10 | -0.00 | 0.21 |
| Algae | 0.16 | 0.05 | 0.27 |
| Structure | -0.08 | -0.19 | 0.04 |
| Predators | -0.08 | -0.19 | 0.03 |
| Mean herbivore density at site | 0.16 | 0.02 | 0.30 |
| Algae: Structure | 0.07 | -0.04 | 0.17 |
| All coefficient estimates are on the log scale and correspond to scaled predictor variables. | | | |

**Sensitivity Analysis 4: Sensitivity to choice of priors**

**Table S25:** Model output for model 1 (bite rates) with priors of slopes set to Normal(0,10)

| **Multilevel hyperparameters: Site (number of levels: 6)** | | | |
|---|---|---|---|
| **Parameter** | **Estimate** | **Lower 95% CI** | **Upper 95% CI** |
| sd(Intercept) | 0.40 | 0.03 | 1.20 |
| **Regression Coefficients** | | | |
| Intercept | 0.00 | -0.43 | 0.43 |
| Group size | 0.49 | 0.32 | 0.66 |
| Aggression rate | -0.28 | -0.45 | -0.12 |
| Algae | 0.12 | -0.10 | 0.34 |
| Structure | -0.08 | -0.28 | 0.13 |
| Mean group size at site | -0.24 | -0.70 | 0.20 |
| All coefficient estimates are for scaled outcome (bite rate) and predictor variables. | | | |

**Table S26:** Model output for model 2 (aggression rates) with priors of slopes set to Normal(0,10)

| **Multilevel hyperparameters: Site (number of levels: 6)** | | | |
|---|---|---|---|
| **Parameter** | **Estimate** | **Lower 95% CI** | **Upper 95% CI** |
| sd(Intercept) | 0.26 | 0.01 | 0.88 |
| sd(Hurdle intercept) | 0.78 | 0.04 | 2.27 |
| **Regression Coefficients : Hurdle** | | | |
| Intercept | -0.53 | -1.38 | 0.36 |
| Group size | 0.24 | -0.22 | 0.73 |
| Algae | -0.47 | -1.05 | 0.13 |
| Total aggressors | -0.10 | -0.59 | 0.37 |
| Mean group size at site | 0.00 | -0.96 | 0.94 |
| **Regression Coefficients: Gamma** | | | |
| Intercept | -0.29 | -0.65 | 0.01 |
| Group size | -0.14 | -0.30 | 0.03 |
| Algae | 0.08 | -0.14 | 0.29 |
| Total aggressors | 0.13 | -0.06 | 0.32 |
| Mean group size at site | 0.02 | -0.29 | 0.37 |
| All coefficients estimates are for scaled predictor variables. The coefficients for the hurdle model are on the logit scale, and coefficients for the gamma part are on the log scale | | | |

**Table S27:** Model output for model 3 (proportion grouping) with priors of slopes set to Normal(0,10)

| **Multilevel hyperparameters: Site (number of levels: 15)** | | | |
|---|---|---|---|
| **Parameter** | **Estimate** | **Lower 95% CI** | **Upper 95% CI** |
| sd(Intercept) | 0.22 | 0.01 | 0.58 |
| **Regression Coefficients** | | | |
| Intercept | 1.16 | 0.91 | 1.41 |
| Herbivores | 0.77 | 0.45 | 1.10 |
| Territorials | 0.34 | 0.11 | 0.58 |
| Algae | 0.05 | -0.19 | 0.28 |
| Structure | -0.18 | -0.40 | 0.03 |
| Predators | 0.00 | -0.24 | 0.29 |
| Mean herbivore density at site | -0.05 | -0.34 | 0.24 |
| All coefficient estimates are on the logit scale and correspond to scaled predictor variables. | | | |

**Table S28:** Model output for model 4 (mean group sizes) with priors of slopes set to Normal(0,10)

| **Multilevel hyperparameters: Site (number of levels: 15)** | | | |
|---|---|---|---|
| **Parameter** | **Estimate** | **Lower 95% CI** | **Upper 95% CI** |
| sd(Intercept) | 0.09 | 0.00 | 0.25 |
| **Regression Coefficients** | | | |
| Intercept | 2.04 | 1.93 | 2.16 |
| Herbivores | 0.54 | 0.42 | 0.67 |
| Territorials | 0.10 | -0.01 | 0.21 |
| Algae | 0.14 | 0.03 | 0.25 |
| Structure | -0.08 | -0.19 | 0.04 |
| Predators | -0.09 | -0.19 | 0.02 |
| Mean herbivore density at site | 0.17 | 0.03 | 0.31 |
| All coefficient estimates are on the log scale and correspond to scaled predictor variables. | | | |
